# Animal models of epilepsy: legacies and new directions

**DOI:** 10.1101/013136

**Authors:** Brian P. Grone, Scott C. Baraban

## Abstract

Human epilepsies encompass a wide variety of clinical, behavioral and electrical manifestations. Correspondingly, studies of this disease in nonhuman animals have brought forward an equally wide array of animal models, i.e. species and acute or chronic seizure induction protocols. Epilepsy research has a long history of comparative anatomical and physiological studies on a range of mostly mammalian species. Nonetheless, a relatively limited number of rodent models emerged as the primary choices for most epilepsy-related investigations. In many cases these animal models are selected based on convenience or tradition, though technical or experimental rationale does, and should, factor into these decisions. More complex mammalian brains and, especially, genetic model organisms including zebrafish have been studied less but offer significant advantages that are being widely recognized.

## Introduction

First and foremost, there is no single animal model of epilepsy that fully represents this disease. Perhaps this is not surprising given the range of conditions subsumed under the term “epilepsy”, each with distinct acquired or genetic origins and diverse behavioral manifestations, electrographic signatures, pharmacological profiles, and histopathologies. An animal model of epilepsy generally refers to a particular experimental species and an induced or inherited propensity for developing seizures. Like all disease models these animals should recapitulate (i) the causal mechanism(s) underlying the disease in humans (construct validity), (ii) phenotypic features of the human condition (face validity) and (iii) treatment responses seen clinically (predictive validity). In a hypothetical sense, the perfect model would satisfy all three criteria, i.e. have similar etiology as a human form, exhibit the same physiological, behavioral or genetic phenotypes, and respond to the same therapies^1^. In a practical sense this goal has remained elusive since the first laboratory demonstration of electroshock-induced seizures. For a comprehensive overview of animal models of epilepsy we refer the reader to the many texts that describe and characterize different standard models^2–12^. What follows here is a discussion of issues related to the history of animal model development, and some of the emerging approaches that allow epilepsy to be modeled in a wide variety of species.

Human epilepsies are most often defined as exhibiting “recurrent self-sustained paroxysmal disorders of brain function characterized by excessive discharge of cerebral neurons”^10^. Therefore, animal models of epilepsy seek to recapitulate and elucidate these abnormal electrographic discharges as well as many of the underlying neuroanatomical, biochemical^13–15^, and genetic^16–20^ factors that lead to them. In many animal models, electrographic seizures, verified via an electroencephalogram (EEG), are also closely associated with observable and quantifiable convulsive behaviors. Seizures can be observed following acute insults or genetic manipulations in nearly every animal with a nervous system, so the potential range of models is vast. The fundamentally shared propensity of animals for developing seizures was appreciated as early as 1869 when John Hughlings Jackson called for greater attention to comparative physiology in studying spontaneously arising seizures in dogs^21^. Pioneering studies by his colleague David Ferrier on rabbits, guinea pigs, cats, and dogs soon demonstrated that direct electrical stimulation of cortex in these mammals generates clonic seizure events that resemble human epilepsy^22^. It is important to clarify that electrically induced seizures are acute events rather than chronic models of spontaneous recurrent epileptic seizures.

Such electrically induced seizures were used in cats to test anticonvulsant drugs during the 1930s and beyond^23^. When intracellular and extracellular recording techniques became prevalent in the late 1970s, these acute models proved useful in elucidating electrical components underlying seizure events – paroxysmal depolarization shift (PDS) in particular^10^. Generally, acute seizure models are best when used to model the one-third of human epilepsies classified as acquired. Complementing this work, induced or acquired seizure protocols were developed, e.g. maximal elcectroshock (MES) which models generalized tonic-clonic seizures and acute pentylenetetrazole (PTZ) injection which models clonic seizures. These models were used to discover many of our currently used antiepileptic drugs (AEDs) acting on ion channel or receptor targets^8, 9, 17^.

The history of epilepsy research shows that, more than theoretical perfection, experimental tractability often determines which approaches are adopted and pursued. Although epilepsy investigators made extensive use of dogs, non-human primates, and especially cats ^24, 25^ before the 1980s, rats and mice with acquired forms of epilepsy have become, by far, the most common type of animal model. Widespread and historical use of rodents often makes them the default animal for these types of experiments. While there are many continuing advantages to these rodent models, and a rich literature for comparison, it is wise in any experimental model choice to match the advantages of the model with the research questions posed.

## Comparative Animal Models

When choosing an animal model, it should be kept in mind that all animal studies are in essence comparative. A comparative approach can reveal aspects of epileptogenesis that are widespread in animals (Figure 1), and thus likely to be clinically important in humans, as well as deeper understanding of specific processes accessible in particular species. August Krogh asserted that “for such a large number of problems there will be some animal of choice, or a few such animals, on which it can be most conveniently studied.” ^26^. Accepting this, two thorny problems remain: what constitutes an important biological “problem” in the epilepsy field? And, of course, which experimental animal is the best suited to address said problem?

Historically, a range of species has been convenient for particular purposes within the study of epilepsy. Each species has particular similarities to, and differences from, humans that influence the types of studies for which they are best suited. As mentioned above, cats, dogs, and non-human primates were commonly used to study experimentally induced seizures throughout the first decades of epilepsy research^6^. For example, alumina cream was discovered serendipitously to generate generalized electrographic seizures when applied to the brains of monkeys^27^. Their large gyrencephalic mammalian brains remain more similar to our own than are those of many simpler model organisms. Longitudinal datasets from captive baboons revealing epidemiological^28^ and neuroanatomical^29^ features of epilepsy have the potential to contribute to understanding genetic bases of seizures in animals with brains and genomes very similar to our own. Similarly, dogs with spontaneous naturally occurring seizures, remain valuable for the development of EEG-based seizure detection algorithms^30^ and evaluation of novel therapies^31^. Canine epilepsies have face validity, mimicking the clinical manifestation of the nearly one-third of human epilepsies classified as idiopathic. Yet it should be cautioned that the unknown origins of epilepsy in heterogeneous collections of dogs complicate interpretation of these studies^32^. On-going studies of drug efficacy in canine models featuring chronic seizures are clearly warranted, and have continued long after most work on cats and primates came to a halt.

Less commonly used species also have much to offer. Invertebrates, including a nudibranch mollusc, *Tritonia diomedea*, are useful in studying basic neural mechanisms underlying differential responses to, for example, brain injury^33^, a common cause of epilepsy. Simple preparations like the three-neuron-type oscillator that generates locomotor rhythms in *T. diomedea* offer a high degree of accessibility for surgeries, neural recording, and behavior. This experimental tractability makes it a potentially convenient model to answer questions about recurrent excitatory circuits, a long-standing theoretical basis for seizure generation in reorganized epileptic hippocampal circuits^34^. *Xenopus* tadpoles are a classical model for neurodevelopmental studies, and were recently used to study PTZ-induced seizures. These studies revealed biochemical mechanisms of resistance to subsequent seizures^35^. Although PTZ treatment in tadpoles in some ways falls short as a preclinical model of human epilepsy, given that seizure threshold increases after a treatment rather than decreasing spontaneously, the differences may be illustrative for mechanisms controlling seizure sensitivity. Turtles, which are highly resistant to hypoxia, allow studies of intact brains in vitro^36^ as well as unusual resistance to hyperexcitability and cell death^37^. In birds, classic embryological transplants allow creation of chimeric brains for the study of epilepsy foci^38, 39^. More recently, sea lions with epilepsy resulting from ingestion of domoic acid have been shown to develop hippocampal pathology resembling what is seen in humans with temporal lobe epilepsy^40^. Broadly based study of epilepsy mechanisms in these species and others, though challenging and perhaps not always convenient, can complement ongoing work in traditional animal models of epilepsy.

Though mutagenesis and transgenesis for epilepsy studies is most commonly applied to mice, it is also possible in other species, for example rabbits^41, 42^, pigs^43^, and primates^44^. For areas such as long-term evaluation of slowly maturing neurons derived from human embryonic or induced pluripotent stem cell sources, transgenesis in pigs or rabbits may be necessary.

## On Epilepsy in Rodents

Rodents, in particular the Norway rat (*Rattus norvegicus*) and the house mouse (*Mus musculus*) are important species for epilepsy research as well as neurodevelopment and biomedicine more broadly. The ascendance of these two species may be linked to their small size, docility, and rapid breeding in captivity. A substantial literature exists describing cell firing properties, synaptic function, neurotransmitter actions, etc. in “normal” lab mice and rats. Gene sequence and expression data for model organisms, emerging in the 1980s and beyond, added to a significant literature using these acquired rodent seizure models to identify many of the neuroanatomical consequences of seizure activity. Studies of epileptic rodents uncovered many of the deficits we now associate with an epileptic brain.

Rodent models of acquired epilepsy have contributed greatly to our understanding of the relationship between surface EEG events and the underlying firing activity of individual neurons. Seizures in rodents were reported with focal application of cobalt, tungstic acid, acetylcholine, strychnine, or picrotoxin^9, 10^. To progress from studying acutely generated seizure events to studying a chronic phenotype more closely resembling epilepsy (i.e. spontaneous recurrent seizures) new approaches emerged in the 1970s. Chronic repetitive electrical (or chemical) stimulation of the brain generates a phenomenon known as kindling, in which the threshold for electrically stimulated seizures decreases, and spontaneous seizures can develop^45^. Kindled animals are considered a model of complex partial seizures with secondary generalization and have proven useful for establishing seizure thresholds and how they may, or may not, respond to therapeutic treatments. However, the relevance of kindling models to specific forms of human epilepsy remains controversial^46^. Systemic pilocarpine or kainic acid administration similarly can evoke acute episodes of status epilepticus and, with time, recurrent spontaneous seizures with clear behavioral manifestations. These models enjoy widespread use and are easy to implement. Owing to the neuronal damage and synaptic reorganization that these models generate, i.e. sprouting of granule cell excitatory axon collaterals (mossy fibers), they are often referred to as animal models of temporal lobe epilepsy. These models are well suited to studying mechanisms and biomarkers of epileptogenesis, or validating novel drug discoveries that may emerge from high-throughput screening. It has, however, been cautioned that targeting these models (developed in otherwise healthy rodents) for continued therapeutic development may reveal only treatments closely related to known ones^8^.

While it may often be taken for granted that effects found in mice closely resemble those present in humans, counter-examples exist, due to differences in gene expression, protein function, and gene network participation^47^. And within any species, differences in genetic background exist among heterogeneous populations and inbred strains^48, 49^. These differences should be considered in choosing models. For example, some strains of mice are more resistant to kainic acid than others^50^. Similarly, even between closely related species such as rats and mice there are differences in degree and function of adult neurogenesis^51^, a potentially critical aspect of the epileptogenic process as seizures evoke a strong neurogenic response in the hippocampal subventricular zone^52^. Furthermore, susceptibility of different strains of mice to seizures does not always correlate with physiological substrates thought to underlie epilepsy, including neuron loss and mossy fiber sprouting^53^. These differences do not necessarily negate the relevance of chemoconvulsant rodent models, as humans with acquired forms of epilepsy also do not uniformly show neuron loss and mossy fiber sprouting^54^. Nonetheless, from a scientific point of view, rodent models should be chosen when appropriate for their outstanding features rather than due to the sheer inertia of the research enterprise that has been built upon their backs.

The identification of spontaneous, and later induced, genetic epilepsies in mice has proven invaluable for modeling human epilepsies. In 1957 a family of mice at Jackson Laboratories were found to have an abnormal, wobbly gait^55^. These *Tottering* mice were soon identified as having alleles of the same gene on chromosome 8 that was also mutated in *Leaner* mice^56^. They exhibited absence seizures and intermittent focal seizures, consistent with human phenotypes^57^. Subsequently, it was recognized that mutant mice with behavioral and electrophysiological seizures could serve as models for human disease, although the molecular basis for their symptoms were unknown^58^ until it was shown that *Tottering* and *Leaner* mice carry mutations in an alpha1a calcium channel^59^. Mice with spike-wave discharge and absence-like seizures were subsequently found to have mutations in subunits of voltage-gated calcium channels, suggesting a core set of molecular mechanisms that could model a set of related human seizure disorders^60, 61^. With advancements in gene editing techniques, particularly homologous recombination, a variety of additional mice have now been identified as “epileptic” (e.g., exhibiting spontaneous recurrent seizures) or “seizure susceptible” (e.g., exhibiting a reduced threshold for acute induction of seizures). In many cases, transgenic mice now exist to mimic specific human conditions such as Type I Lissencephaly^62^, Tuberous Sclerosis Complex^63^ or Dravet syndrome^64^. Investigations using these mouse models facilitated identification of underlying circuit deficits contributing to a hyperexcitable (or epileptic) conditions, the advancement of therapeutic interventions such as rapamycin designed to target specific signaling pathways associated with the disease^65^ or the potential clonazepam-mediated rescue of autism-related co-morbidities^66^. In contrast to mice, rats have so far proven more difficult to use in modeling genetic forms of epilepsy. Rats exhibiting heritable absence seizure episodes have been identified and mimic some aspects of human absence epilepsy, but the underlying genetic mutation responsible for epilepsy in these animals remains a mystery^67^. For some types of experiments, e.g. evaluating epilepsy co-morbidities in a series of complex behavioral tasks, transgene approaches in rats^68^ offer distinct advantages.

## The Rise of Genetic Model Organisms

For many basic neuroscience laboratories, *Drosophila melanogaster*, the fruit fly or vinegar fly, is a model organism par excellence. Genetic modifications are rapid and relatively simple in Drosophila, and many epilepsy genes are conserved (see Table 1). One of the first seizure mutations found in Drosophila, shibire, affects a molecular motor protein, dynamin, which has recently been linked to human epilepsy^69^. Recent work on Drosophila voltage-sensitive sodium channels exemplifies how epilepsies may be modeled this small species whose brains contain approximately 1,000,000-fold fewer neurons than our own. The Drosophila neural sodium channel encoded by *para* shares highly conserved structure and 43% pairwise identity with the sodium channel 1a gene (*SCN1A*) mutated in most cases of Dravet syndrome^70^. As such, specific nucleotide modifications similar to mutations found in humans^71^ can be introduced. Though knock-ins are also possible in mice, with over 300 different mutations identified in *SCN1a* alone^72^, generating mutant mice for each one is a daunting task. Other genes involved in human epilepsies were also identified first in flies by behavioral screening and genetic mapping, i.e. *Shaker*^73^. Subsequent work showed that the homologous Kv1.1 channel is responsible for epilepsies in mice^74^ and humans^75^.

Abnormalities in many different genes (estimated at 1000^76^) can lead to genetic epilepsies, and these are now being identified and characterized^77^, though many remain to be modeled in experimental animals. The large number and diversity of these genes suggests that high throughput may be a decisive advantage in model development, and Drosophila’s small size, low cost, and rapid transgenesis allow probing of many variants. On the other hand, the tough carapace of Drosophila means that drug treatments are not convenient and, because flies are not vertebrates, limitations on interpreting behavior or recording electrographic events in Drosophila brain remain. Other invertebrates, e.g. *C. elegans*, have been described as exhibiting seizure-like phenotypes^78^ but are only distantly related to humans. *C. elegans*, for example, lacks voltage-gated sodium channels^79^ and has a “nerve ring” brain that includes only 180 neurons. Therefore, invertebrate results should be interpreted with caution when considering the more complex human epileptic circuits we seek to model and understand. Nevertheless, genes first discovered in *C. elegans* and Drosophila have proven extraordinarily valuable in understanding basic mechanisms of neural transmission in vertebrates.

In contrast to flies and worms, zebrafish (*Danio rerio*) is a vertebrate genetic model organism with tremendous potential for modeling acute seizures and genetic epilepsies. Zebrafish, small striped minnows from the Indian subcontinent, were introduced to the West early in the 20^th^ century. By the 1930s they were appreciated as a suitable species for developmental biology studies: *“Eggs in considerable number can be obtained every day of the year. The egg is transparent, has a small yolk, and can be held in any desired position without trouble. Furthermore, the development is unusually rapid*”^80^. During the 1960s, George Streisinger appreciated these logistical advantages and surmised that zebrafish would be an excellent model for molecular genetics. This insight proved lucid, as many studies that could previously only have been performed in Drosophila became feasible in a vertebrate that shares 75% of human disease genes^81^. Physiologically and behaviorally, acute seizures induced in wild type zebrafish by 4-aminopyridine and pentylenetetrazole^82^ or heat^83^ closely resemble those induced in mammals. Indeed, a recent mathematical analysis of electrical seizure events in mice, zebrafish and humans concluded that the underlying rules governing initiation and termination were universal^84^.

Although zebrafish are useful to study acutely evoked seizure events, their true advantage lies in modeling genetic epilepsies. Early genetic screens pioneered by Christiane Nusslein-Vollhard identified zebrafish with motor deficits and uncovered an array of mutations in potential epilepsy-linked genes^87, 88^. Building on this work and using zebrafish mutants emerging from such forward-genetic ENU-based screens, our laboratory identified and characterized the first mutant zebrafish models of epilepsy including Ube3a^89^ (Angelman’s syndrome), Scn1a^90^ (Dravet syndrome), and Ocrl1^91^ (Lowe’s syndrome). In zebrafish, field potential recording of synchronized neural activity can readily be performed in series with behavioral tracking experiments allowing unequivocal confirmation of epilepsy in these models^82, 83, 89, 90, 92–95^. High-throughput mutagenesis efforts aiming to mutate every gene in zebrafish are well underway, and provide an unparalleled resource for characterizing additional epilepsy phenotypes resulting from clinically relevant genes^96, 97^. For generating specific mutations or gene knockdowns, the standard loss-of-function approach in zebrafish has until recently been injection of morpholinos i.e., modified oligonucleotides that bind to and block translation of endogenous mRNA. This approach is effective during early development and allowed rapid modeling of epilepsy genes including kcnq^94^, lgi1^98^ and *chd2*^99^. However, drawbacks include the transience of morpholinos, variable knockdown, and off-target effects^100^. By adapting the CRISPR/Cas9 antiviral system from bacteria, recent work further increased the ease of straightforward targeting of specific DNA sequences in a broad range of species^101^ allowing rapid knockout as well as knockin of specific alleles in zebrafish. CRISPR/Cas9, unlike Zinc Finger Nucleases and TALENs, does not require multiple lengthy cloning steps to generate engineered genes encoding proteins that interact with and cleave DNA. Relying instead on complementary DNA-RNA binding the CRISPR/Cas9 system can target virtually any non-repetitive genomic region. These newer gene editing techniques open up the possibility that mutant zebrafish lines can be generated for all known, and yet-to-be-discovered, human epilepsy genes.

Although the miniscule size of larval zebrafish currently prevents the simultaneous recording of behavior and EEG, zebrafish offer some unique advantages for epilepsy research. For example, zebrafish membranes are permeable to drugs placed in the bathing medium. The ease with which drugs can be delivered to freely behaving zebrafish and the sensitivity of locomotion and electrophysiology assays for seizure activity resulted in a novel moderate- to high-throughput platform for the discovery of new AEDs. Recent work from our laboratory used a zebrafish line carrying a mutation in the *scn1lab* gene with 78% homology to the human *SCN1A* gene that has been associated with Dravet syndrome. Screening a re-purposed drug library of more than 300 compounds led to the discovery of clemizole^90^, a drug that had never been tested against seizures in rodents and likely never would have been. Rapid targeted mutagenesis and drug screening provide obvious advantages for a personalized form of drug discovery in genetic epilepsies. Furthermore, zebrafish larvae, particularly the *nacre* strain, are transparent allowing in vivo imaging of fluorescently labeled cells, genetically encoded calcium indicators, and fluorophore-tagged proteins ^102, 103^. Combined with advanced techniques for brainwide optical monitoring of patterned neural activity – light sheet and spinning disk confocal microscopy – additional avenues for studying epilepsy in this simple vertebrate species are evolving.

## Conclusion

A diverse toolkit facilitates modeling epilepsy in animals. Our understanding of this disease process has benefitted greatly from established acute and chronic models. However, pairing modern neuroscience strategies with comparisons across diverse models offers a more focused question-specific approach to using models. It is important to recognize seminal insights and treatments that arose from traditional rodent models, but at the same time, consider the limitations of these rodent models, and where new technologies are emerging to facilitate a broader approach to modeling. Given these considerations, it makes sense for future epilepsy investigators to have an open mind and embrace a wide range of animal models – flies, zebrafish, mice, pigs, sea lions, rabbits, etc. – in their pursuit of a better overall understanding of the epileptogenic process and the development or discovery of novel therapies for this common neurological disorder.

**Table 1.**
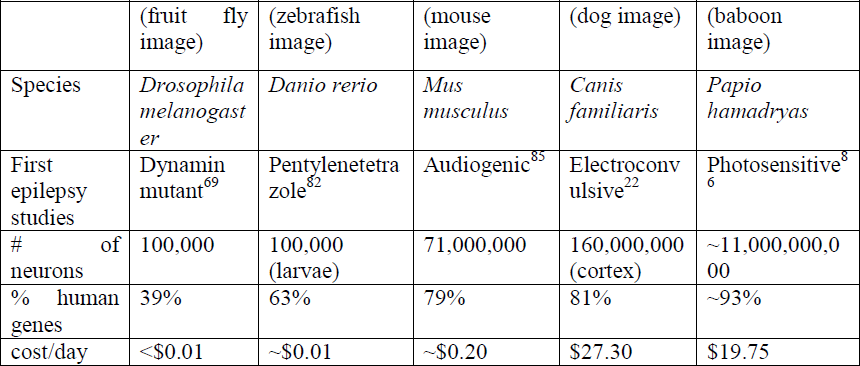
Epilepsy model organisms span the range from simple to complex species. Costs according to UCSF per diem rates

|  |  |  |  |  |  |
| --- | --- | --- | --- | --- | --- |
|  | (fruit fly image) | (zebrafish image) | (mouse image) | (dog image) | (baboon image) |
| Species | *Drosophila melanogaster* | *Danio rerio* | *Mus musculus* | *Canis familiaris* | *Papio hamadryas* |
| First epilepsy studies | Dynamin mutant<sup>69</sup> | Pentylentetrazole<sup>82</sup> | Audiogenic<sup>85</sup> | Electroconvulsive<sup>22</sup> | Photosensitive<sup>86</sup> |
| # of neurons | 100,000 | 100,000 (larvae) | 71,000,000 | 160,000,000 (cortex) | ~11,000,000,000 |
| % human genes | 39% | 63% | 79% | 81% | ~93% |
| cost/day | <\$0.01 | ~\$0.01 | ~\$0.20 | \$27.30 | \$19.75 |

## Acknowledgements

S.C.B. is supported by funds from the NIH, Citizens United for Research in Epilepsy, Dravet Syndrome Foundation and California Institute for Regenerative Medicine. B.P.G. is supported, in part, by funds from the Lennox-Gastaut Syndrome foundation. We here acknowledge the difficult task of summarizing, categorizing and comparing the many experimental models of epilepsy that exist and apologize to those whose work could not fit into the brief confines of this manuscript. The authors would also like to thank Robert Hunt and MacKenzie Howard for insightful discussions and advice regarding the writing of this Perspective.

